# Identifying myocardial regions perfused by coronary arteries through detailed human microvasculature data

**DOI:** 10.1101/2025.05.12.653398

**Authors:** Giovanni Montino Pelagi, Jay A. Mackenzie, Ed Van Bavel, Giovanni Valbusa, Christian Vergara, Nicholas A. Hill

## Abstract

**Purpose:** clinical imaging can resolve the main coronary arteries but not the smaller side branches that penetrate the heartwall. However, precise association between the main coronary branches and the myocardial mass they perfuse is crucial to achieve a correct description of haemodynamics from the large arteries to the cardiac tissue. In this work, we use ex-vivo detailed morphometric data of human coronary microcirculation to build and validate a tool for a personalized coronary-myocardium association, and we use it in a multiscale computational model of cardiac perfusion.

**Methods:** from the digitalized dataset of an entire human coronary microcirculation, vascular beds associated to single branches are extracted and analysed to infer patterns in epicardial branching. 3D segmentations of the coronaries with and without this information are used to generate two different myocardial subdivisions, which are compared to the one obtained from the microcirculation data. The impact on haemodynamics is assessed through computational simulations.

**Results:** epicardial arteries exhibit characteristic patterns of transverse branching, with branching angles ≃ 90° and rate of branching, with respect to the distance along the vessel, depending on the core diameter. The addition of transverse outflows to the segmentations greatly increases accuracy in the myocardial subdivision, allowing discrimination between mass perfused by the proximal and distal arterial segments. Perfusion simulations including transverse outflows show more homogeneous blood flow across the myocardium, consistently with experimental findings.

**Conclusions:** the inclusion of transverse outflows in 3D coronary segmentations is essential to correctly capture the coronary-myocardium association and the distribution of myocardial blood flow.

## 1. Introducation

The coronary arterial circulation distributes blood throughout the cardiac tissues and consists of a tree-like structure originating from the aortic root and branching down to the capillaries, embedded within the myocardial fibrils. In pathological conditions such as Coronary Artery Disease (CAD), understanding how a lesion in a given major coronary artery translates into cardiac ischaemia in the myocardium and quantifying which regions are affected are still problems that motivate efforts to obtain detailed vascular data in research settings. Such data have been obtained from arterial casts in pigs [1, 2, 3] and recently in humans [4], highlighting a specific branching pattern: the core of the larger coronaries runs on the epicardium with a high number of much smaller transverse vessels branching off and penetrating the myocardium.

This suggests that the myocardial territories perfused by each of the epicardial branches are elongated in shape and follow the path of their associated feeder arteries, reflecting their anatomical variability. The clinical relevance of a correct quantification of the Mass at ischaemic Risk (MaR) in patients affected by CAD [5] prompted the development of image-based, patient specific methods for the association between coronary feeding arteries and myocardial mass [6, 7, 8, 9]. Some of these studies also found that the ischaemic burden is correlated to both the MaR and the functional relevance of the lesion.

A very promising to quantify the effects of epicardial lesions on cardiac perfusion is to use of computational models coupling the haemodynamics in the large arteries with regional perfusion. Some of these works included detailed morphometric data to provide an accurate spatial link between each coronary branch and the myocardial mass they perfuse [10, 11]. These data, however, are not available for living subjects, where the anatomy is provided by only *in vivo* medical imaging. To address this issue, in other works [12, 13, 14] the myocardium is divided into perfusion regions starting from the outlets of a coronary tree segmented from Computed Tomography (CT) or Magnetic Resonance (MRI) images that have a minimum resolution ≃ 0.5 mm. A critical point is that the majority − of the transverse vessels have a diameter in the range *d* = 0.3 − 0.7 mm and are, therefore, missed in the coronary segmentation. The myocardial subdivision proposed in these works, therefore, does not take into account the presence of these additional vessels, leading to potentially inaccurate associations between major coronary branches and their corresponding perfusion territory. Other works [15] exploited morphometric pig data [2] to generate realistic synthetic vasculature including transverse vessels, but a comprehensive study based on human data and providing quantitative validation is still lacking. How to perform an accurate myocardial subdivision for any given patient starting from non-invasive imaging remains, therefore, an open issue in the field.

In our work, we push further the analysis of the detailed human vascular data reported in [4] with the purpose of establishing *quantitative* relationships between epicardial branches, transverse vessel topology and myocardial mass. We then integrate this information into a new method for a personalized subdivision of the myocardium into perfusion territories, with an anatomically accurate association between cardiac mass and either proximal, medial or distal region of each feeding artery, and we validate it against the *ground truth* perfusion regions extracted from the detailed vascular data. We finally perform perfusion simulations (using the computational model developed in [16]) to explore the impact of our strategy on coronary haemodynamics and myocardial blood flow distribution.

To the best of our knowledge, this is the first time that detailed human vascular data have been integrated in a tool to achieve accurate coronary-myocardium association. This tool has the important feature of being designed for its application to 3D geometries of patients derived from *in vivo* images, enabling for the first time the use of fully 3D computational simulations of coronary and myocardial blood flow. It is also the first time that such tool has been validated against *ground truth* perfusion regions coming directly from detailed vascular data. We believe this is a crucial step towards optimal MaR quantification in CAD patients and towards more accurate prediction of myocardial blood flow distribution using computational simulations.

## 2. Branching patterns of epicardial arteries

The analysed dataset was obtained from an *ex vivo*, post mortem arterial cast of a 84-year-old female with no clinical history of any major cardiac event; the dataset was first presented and analysed in [4]. The methods used to analyse the dataset are detailed in Section 2.1, alongside the specific aspects we focus on, with the results reported in Section 2.2.

### 2.1. Branching patterns analysis: methods

Detailed vascular data including topology (vessels’ connectivity) and geometry (vessels diameter, length and centrepoint coordinates) were taken directly from source material, whereas the full 3D geometries of the coronary tree and the left ventricular free wall are obtained by us through semi-automated segmentation from the original cryomicrotome images. The segmentation of the coronary tree includes only the main arteries and branches with diameter *d* > 0.8 mm to mimic the most accurate reconstruction achievable from *in vivo* CT images.

From the vascular data and the 3D segmentations we can infer that: the left ventricular mass is 120 g; the circulation type is left-dominated, with the entire left ventricular mass perfused from the left circulation; a total of 10 main branches with epicardial course are identified i.e. the Left Anterior Descending (LAD) artery alongside 3 diagonal branches (*d*_1_ − *d*_3_) and the Left Circumflex (LCX) artery, alongside a small *ramus intermedius* (*r*_*int*_, *d* < 0.75 mm) and 4 marginal branches (*m*_1_ − *m*_4_), iv) two main branches with interseptal course are identified (*s*_1_, *s*_2_). Table 1 reports a summary of the identified epicardial branches, while the segmented 3D geometries are reported in Figure 1a. Note that the *r*_*int*_ is not present due to its small diameter, which is below the chosen threshold.

**Table 1:** Summary of the 10 identified branches with epicardial course, alongside their acronym and the main coronary artery they originate from.

| Branch acronym | Branch name | Main coronary artery |
| --- | --- | --- |
| $d_1$ | First diagonal | LAD |
| $d_2$ | Second diagonal | LAD |
| $d_3$ | Third diagonal | LAD |
| $s_1$ | First interseptal | LAD |
| $s_2$ | Second interseptal | LAD |
| $r_{int}$ | Ramus intermedius | LCX |
| $m_1$ | First marginal | LCX |
| $m_2$ | Second marginal | LCX |
| $m_3$ | Third marginal | LCX |
| $m_4$ | Fourth marginal | LCX |

**Figure 1:**
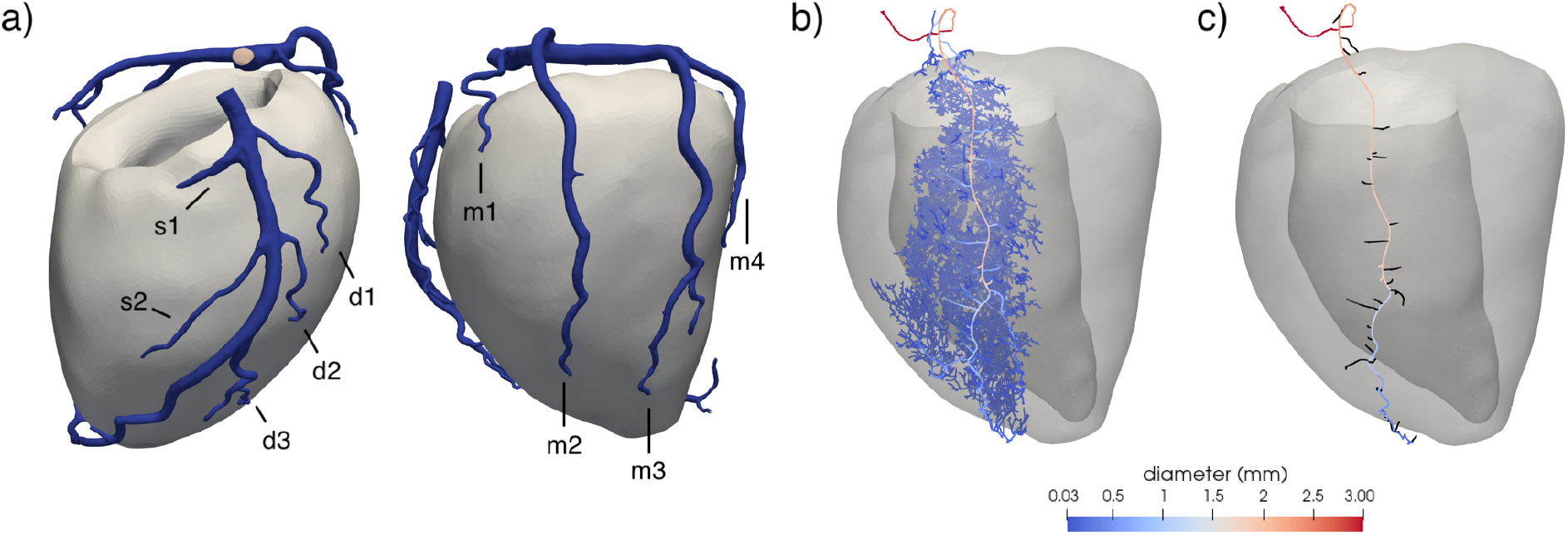
a) 3D segmentations of the left ventricular free wall and coronary tree, where a minimum diameter threshold of *d* = 0.8 mm is used, with indication of the main branches segmented. b) Full isolated vasculature of the second marginal branch (*m*_2_). c) Extracted core of the *m*_2_ branch alongside the first generation of its transverse vessels (in black).

From the entire dataset, vascular subsets associated with each of the main branches (LAD, *d*_1_ − *d*_3_, *s*_1_ − *s*_2_, LCX, *r*_*int*_, *m*_1_ − *m*_4_) are extracted. For each branch, the *core* of the vessel is extracted as the path that runs along the epicardium. This is achieved with a recursive procedure: at each bifurcation point, the vessel running parallel to the epicardial surface is labelled as part of the core, whereas the other vessel(s) (branching close to the perpendicular direction) are labelled as first generation *transverse* vessels. At the distal points where the cores penetrate the myocardium, the criterion used to separate core and transverse segments becomes diameter-based: at each bifurcation point, the vessel with the highest diameter is labelled as core, whereas the lower diameter vessel(s) are labelled as transverse. A graphical representation of the results is shown in Figure 1b-c.

To extract quantitative information on the branching patterns, we define an along-vessel distance *s* following the centerline of the core, such that *s* = 0 corresponds to the starting point of the main artery associated with that branch (LAD or LCX).

### 2.2. Branching patterns analysis: results

In accordance with previous microscopy observations [17], we observe compartmentalized vascular beds that follow the course of the main epicardial branches that they originate from. Figure 2a-b-c reports the cumulative number of transverse vessels for three example branches (*r*_*int*_, *m*_1_ and *d*_1_) as a function of *s*, alongside the core diameter. The results suggest that the branching patterns change along the course of the vessel. In the proximal sections of the epicardial branches, the branching rate (i.e. the number of transverse vessels per unit distance *s*) is constant, with values reported in Table 2. The branches also progressively taper: we report a tapering diameter reduction of 0.19 mm/cm, that is consistent with previous data [18]. In the distal portions, the bifurcation rate increases, with the number of transverse vessels starting to increase at a much faster rate together with a faster drop in vessel diameter; this happens at a core diameter *d*^∗^ ≃ 0.5 mm, consistently for all branches. These results indicate that there is a threshold in diameter at which epicardial arteries start branching with much higher rate, which occurs at the arteries’ distal ends as they penetrate the myocardial mass.

**Table 2:** Results of the branching pattern analysis for all branches: mean and corrected diameters (see eq. (1)), branching rate of transverse vessels and myocardial mass perfused. The *epicardial* rate refers to the epicardial course of the branch, whereas the *deep* rate refers to the distal section, after the branch penetrates the myocardium.

| Branch | $d$ (mm) | $d_{corr}$ (mm) | Branching rate (cm<sup>-1</sup>)<br>Epicardial - Deep | Perfused mass (g) |
| --- | --- | --- | --- | --- |
| $r_{int}$ | 0.52 | 0.52 | 1.63 - 7.29 | 1.10 |
| $m_1$ | 0.94 | 0.94 | 1.26 - 4.72 | 4.13 |
| $m_2$ | 1.40 | 1.40 | 2.38 - 3.56 | 17.30 |
| $m_3$ | 1.54 | 1.22 | 1.66 - 5.21 | 20.00 |
| $m_4$ | 1.08 | 1.08 | 1.54 - 3.14 | 14.60 |
| $d_1$ | 0.84 | 0.84 | 2.57 - 8.88 | 4.80 |
| $d_2$ | 0.66 | 0.66 | 1.49 - 7.66 | 4.30 |
| $d_3$ | 1.29 | 0.82 | 2.56 - 8.79 | 6.40 |
| $s_1$ | 1.30 | 1.30 | 1.75 - 5.56 | 10.80 |
| $s_2$ | 1.10 | 1.10 | 1.39 - 7.21 | 4.50 |

**Figure 2:**
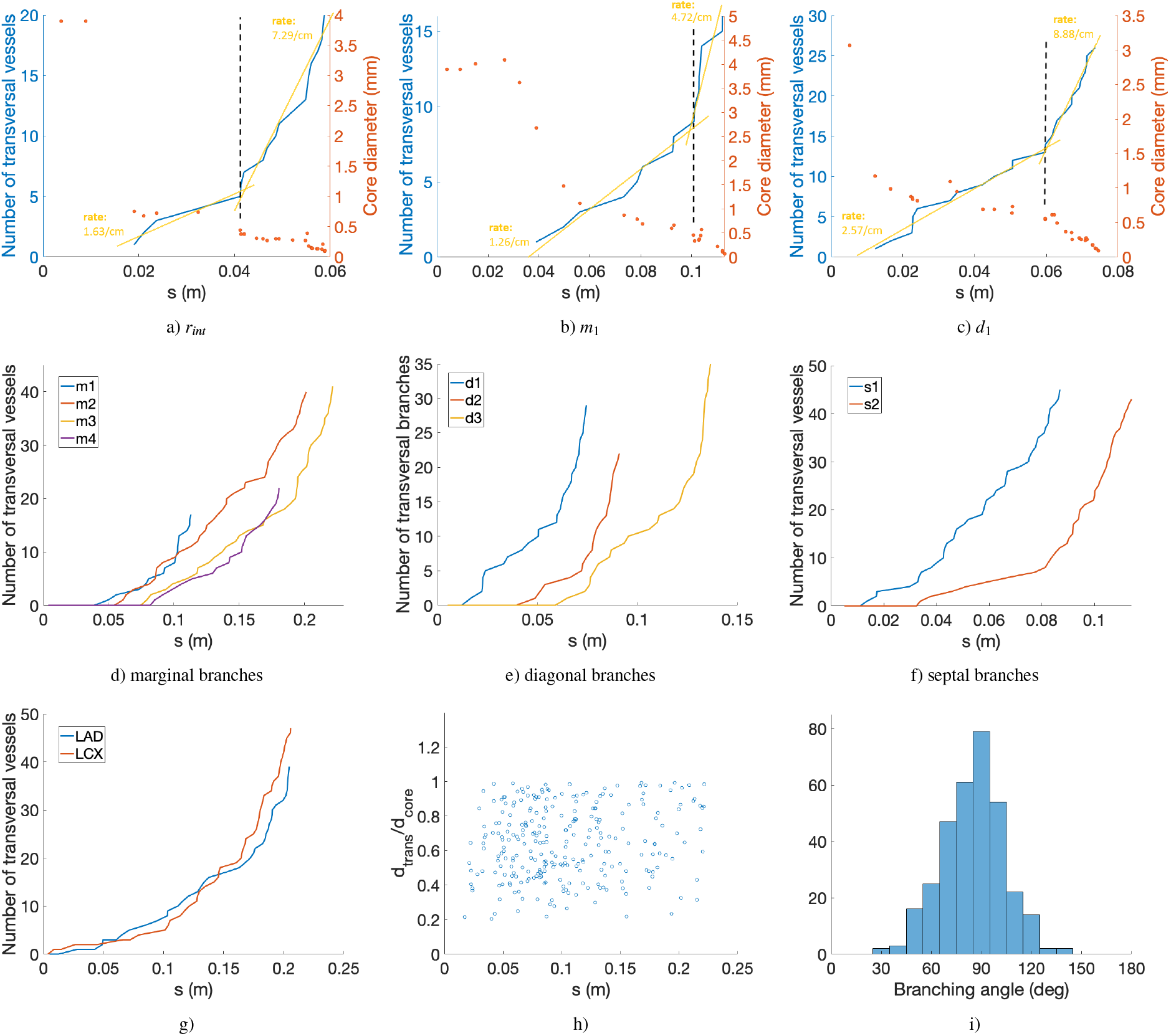
Cumulative branching number for the major coronary branches vs along-vessel distance *s*. a-c) Results for three example branches alongside diameter values of the core vessel structure. d-f) Comparative behaviour for all the branches of the three main types (diagonal, marginal and septal). g) Cumulative number of vessels branching directly from the LAD and LCX vs *s*; h) diameter ratio 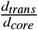 vs *s* for all the branches; i) distribution of the branching angle for the first generation transverse vessels with respect to the core.

Figure 2d-e-f shows the comparative behaviour for all the identified branches. The characteristic piecewise linear shape can be recognized in all the arteries, with diagonal branches exhibiting higher branching rates along their entire course. The only exception regards the vessels branching directly from the cores of the LAD and LCX arteries: as can be seen in Figure 2g, the cumulative number of branches follows a nearly parabolic shape. Also, in the case of the LCX the transverse vessels in between the marginal branches perfuse exclusively the left atrium, while, in the case of the LAD, some ventricle-perfusing vessels start to appear in between the second and third diagonal branch. As can be seen from the scatter plot of Figure 2h, there is high variability in the ratio between the diameter of the transverse vessels and that of the core segment they branch from, with a banded distribution between 0.2 and 1 and no particular dependence on the along-vessel coordinate *s*.

Figure 2i shows the distribution of the branching angles of the transverse vessels with respect to the core. Since the branches can follow a curvilinear trajectory, we first approximate each vessel with a linear, piecewise structure using the change point approach presented in [19]. The angle at each bifurcation is then computed between the direction of the first linear segment of the transverse vessel and the mean direction of the core segments before and after the bifurcation. The distribution is centered at 90^°^ and indicates that the majority of the bifurcations are perpendicular to the core.

The branching rates for all branches are provided in Table 2, together with the diameter and myocardial mass perfused by each branch (see Section 3.1 for details on its computation). For each branch we provide two branching rate values: the *epicardial* rate refers to the epicardial course of the branch, whereas the *deep* rate refers to the distal section, after the branch penetrates the myocardium. These two values of branching rates are computed from the proximal and distal sections of the curves depicted in Figure 2d-e-f, as the slopes of the respective linear interpolants. The diameter *d* represents the mean diameter of the branch and is computed as the weighted average of the diameters of all the segments of the branch’s core (each weight being the segment’s length).

If the branch has bifurcations at the epicardial level, exhibiting two or more large sub-branches running parallel to the epicardium, we introduce a *corrected* diameter *d*_*corr*_ which is computed as:

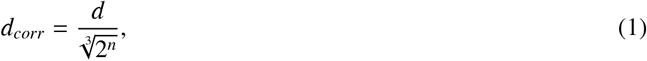

where *n* is the number of consecutive epicardial bifurcation occurring. Eq. (1) is equivalent to Murray’s law in the assumption, at each bifurcation, of two daughter vessels of equal diameter. For all branches without epicardial bifurcations, we have *d* = *d*_*corr*_. This correction is introduced to better differentiate between branches with or without epicardial bifurcations.

The results reported in Table 2 show that larger branches generally show higher branching rates; but these rates are better correlated with *d*_*corr*_ rather than *d*. This means that the presence of bifurcations specifically at the epicardial level results in larger branches without affecting the rate of transverse branching. We also report that the perfused myocardial mass shows a dependence on the actual diameter *d* of the feeding artery also at the level of the single branches, similar to what was already observed at the level of the main arteries [20, 21], although this dependence does not appear to follow strict rules at this level. In this case, epicardial bifurcations are related to a higher mass of perfused tissue.

## 3. Personalized myocardial subdivision strategy

Starting from the analysis reported in Section 2, branching characteristics of the main epicardial coronary branches are inferred and here used to develop a personalized method for accurate subdivision of the myocardial tissue in perfusion regions and correct association between myocardial mass and feeding arteries. In Section 3.1 we describe the new method we propose, while in Section 3.2 we report the results of the proposed subdivision strategy, compared to the outlet-based strategy and the “ground truth” subdivision obtained directly from the detailed vascular data. We also provide a robustness test to analyse how the results are affected by different parameters introduced with the method.

### 3.1. Personalized subdivision strategy: methods

Three different subdivision strategies are employed:

1. *Ground truth*: perfusion regions are obtained directly from the detailed vascular data, which include intramural networks and part of the microcirculation;
2. *Base*: perfusion regions are obtained from the 3D segmentation of the coronary tree, using only the outlets present in the segmentation;
3. *Enhanced*: perfusion regions are obtained from a 3D segmentation of the coronary tree, enhanced with the missing transverse outlets along the main branches.

To obtain the *ground truth* perfusion regions, the complete vascular dataset is divided into branch-specific vascular beds. Each of these beds contain all the vessels that originate from one of the main branches listed in Section 2.1. It is noteworthy that the coronary tree is not, in general, a simple tree with only bifurcations: indeed, confluences and collateral vessels may be present. In the dataset of this study, however, the majority of these confluences involve vessels close to each other and belonging to the vascular bed of the same branch. Only one confluence between different branches *d*_3_ and *m*_2_ was found; in this case, the vessel that bridges the two vascular beds is considered as a junction and is left unassigned.

To achieve a higher precision in myocardial subdivision, we introduce another division of each vascular bed to differentiate between vessels that originate from a more proximal rather than distal portion of the same branch. Specifically, every 3.5 cm of the along-vessel coordinate *s* (see Section 2.1 for its definition) a new subset is introduced. This means that all the vessels originating from the first 3.5 cm of each branch are considered part of the proximal subset of that branch, with additional subsets introduced, as needed, according to the length of the branch. We propose this value of 3.5 cm as a good compromise to retain sensitivity with respect to the proximal-distal distinction without increasing the number of perfusion regions too much. Bifurcations at the epicardial level, as in the case of branches *m*_3_ and *d*_3_, are considered as a change point and each sub-branch is then treated as a separate branch. A graphical representation of the whole subdivision strategy is provided in Figure 3a.

**Figure 3:**
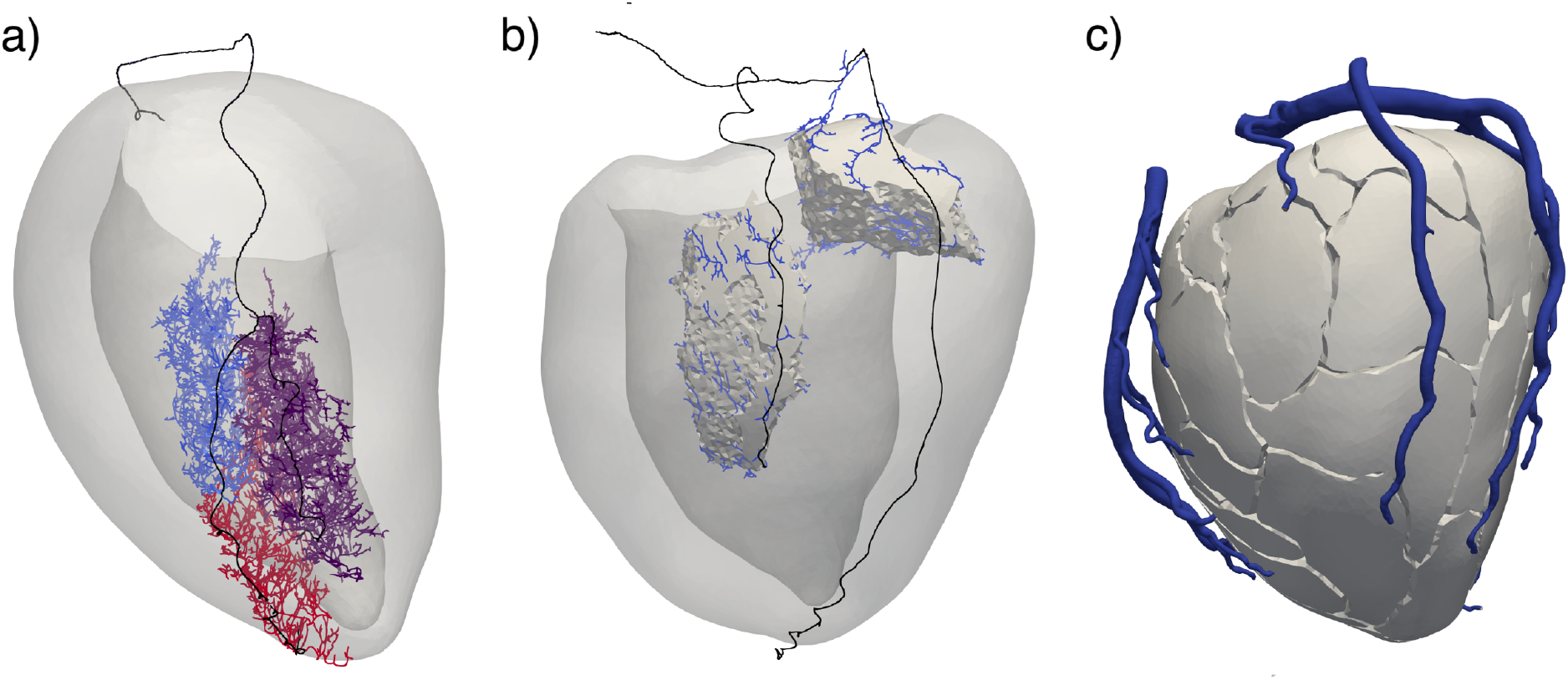
*Ground truth* perfusion regions obtained from detailed vascular data. a) Three different subsets of vasculature originating from the third marginal branch (*m*_3_) distally to a bifurcation at epicardial level: the longest sub-branch is divided into proximal (light blue) and distal (red) parts, with a shorter sub-branch (purple) alongside. b) Wrap surfaces around the points of the ventricular mesh corresponding to two distinct vascular subsets. c) Final subdivision of the ventricular free wall.

For each extracted vascular subset, a boundary surface is wrapped around it using MATLAB AlphaShape function^1^ and the corresponding perfusion region in the ventricle is obtained by finding all the points in the ventricular mesh that lie within such boundary surface. Figure 3b reports a graphical representation of the result for two distinct subsets. The points of the ventricular mesh that are left unassigned after the first step are then assigned to the closest perfusion region, meaning that the wrapping surfaces are slightly inflated to fill the gaps between them. The final subdivision is reported in Figure 3c.

*Base* perfusion regions are obtained starting from the 3D segmentation of the coronary arteries: the centrepoint of each coronary outflow is projected on the myocardial mesh, which is later divided through Voronoi tessellation starting from these points. In this procedure, each point in the ventricular mesh is assigned to the closest projected coronary outflow.

*Enhanced* perfusion regions are still obtained through Voronoi tessellation of the ventricle, but this time the collection of projected points includes also the additional outflows generated on the sides of the major branches. These additional outflows are obtained through the following procedure:

1. The centerline of each epicardial branch is seeded using a given branching rate. Although the results reported in Section 2.2 suggest that such a rate depends on the branch diameter, a constant seeding is used for a first analysis. Two values of 1.4 cm^−1^ and 1 cm^−1^ are used to assess the robustness of the method with respect to this parameter;
2. transverse vessels centerlines are generated connecting each seed to the closest point on the myocardial surface, then the corresponding cylindrical surface is generated. Since there seem to be no clear relation between diameters (see Figure 2h), a diameter ratio of 0.6 is used to be representative of the majority of such vessels;
3. The final 3D surface of the coronary tree with additional transverse outflows is obtained by boolean intersection between the *base* surface and all the transverse vessels.

The procedure and the final 3D coronary surface are reported in Figure 4. From the Voronoi tessellation of the myocardial free wall we directly compute, for each sub-region, the volume and mass assuming a density of 1 g ml^−1^.

**Figure 4:**
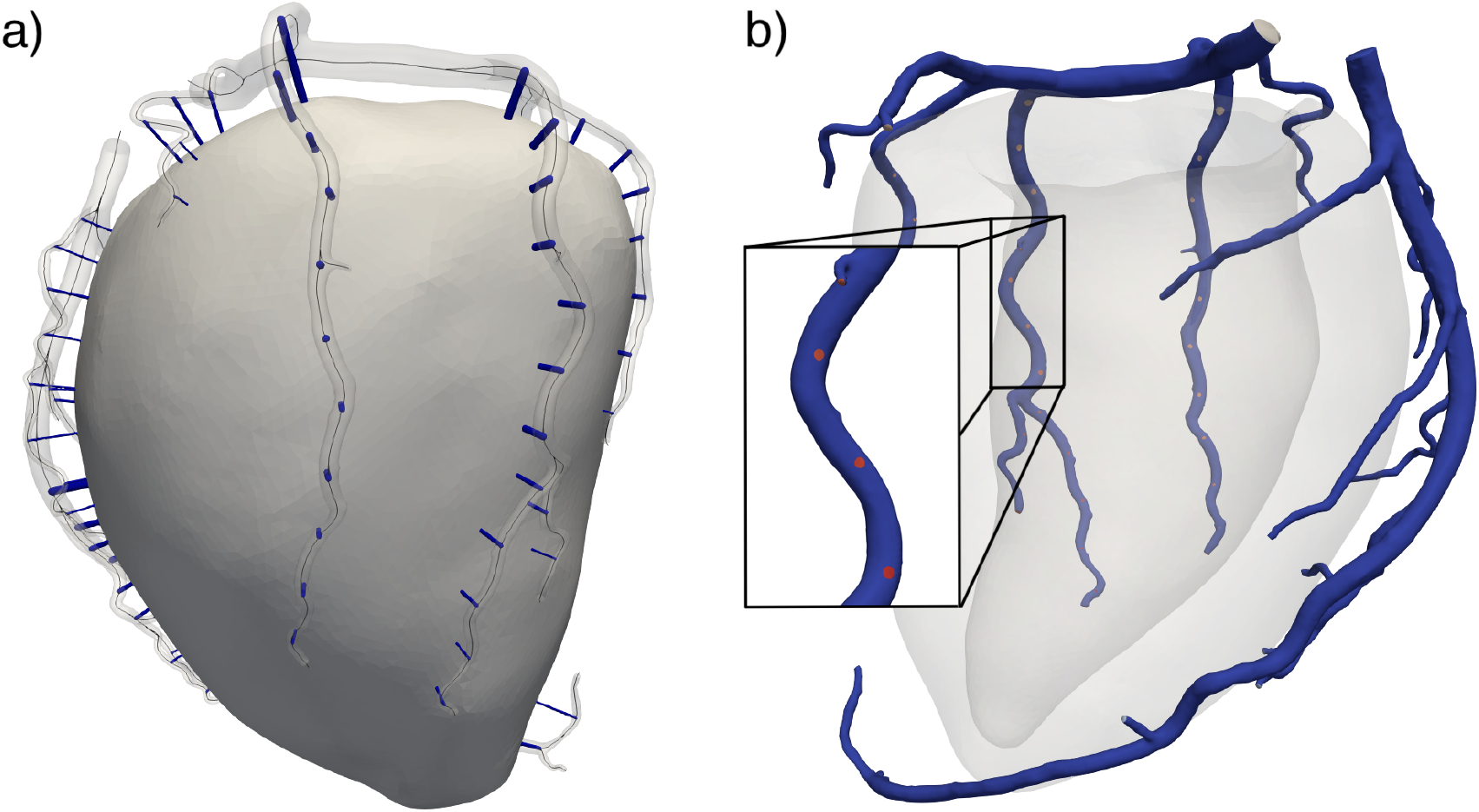
*Enhanced* subdivision strategy: addition of transverse outflows to obtain anatomically accurate perfusion regions. a) transverse vessels generated as cylindrical surfaces connecting seeds on the epicardial coronary branches to their closest point on the myocardial surface, branching rate is 1.4 cm^−1^. b) Final 3D surface of the coronary tree with additional outflows on the wall of the epicardial branches.

### 3.2. Personalized subdivision strategy: results

Figure 5 shows the results of the myocardial subdivision obtained from the *base* and *enhanced* strategies: each ID (characterized by a distinct colour) refers to a single perfusion region associated to a specific coronary outflow. Due to the additional outflows, the *enhanced* strategy results in a much higher number of perfusion regions with a visually different pattern with respect to the *base* strategy. We report a comparison of the union of all regions associated with two example branches (*m*_3_ and *d*_3_, highlighted in gray in Figure 5) with the corresponding *ground truth* regions. The results show that the *enhanced* strategy gives perfusion regions that are characterized by an elongated shape, which is also observed in the vasculature data, and qualitatively much more similar to the corresponding *ground truth* regions than in the *base* case. With the *base* strategy, on the other hand, the proximal regions are likely to be attributed to different branches, leading to mistakes in the artery-myocardium association, as can be seen in particular for branch *m*_3_ in Figure 5.

**Figure 5:**
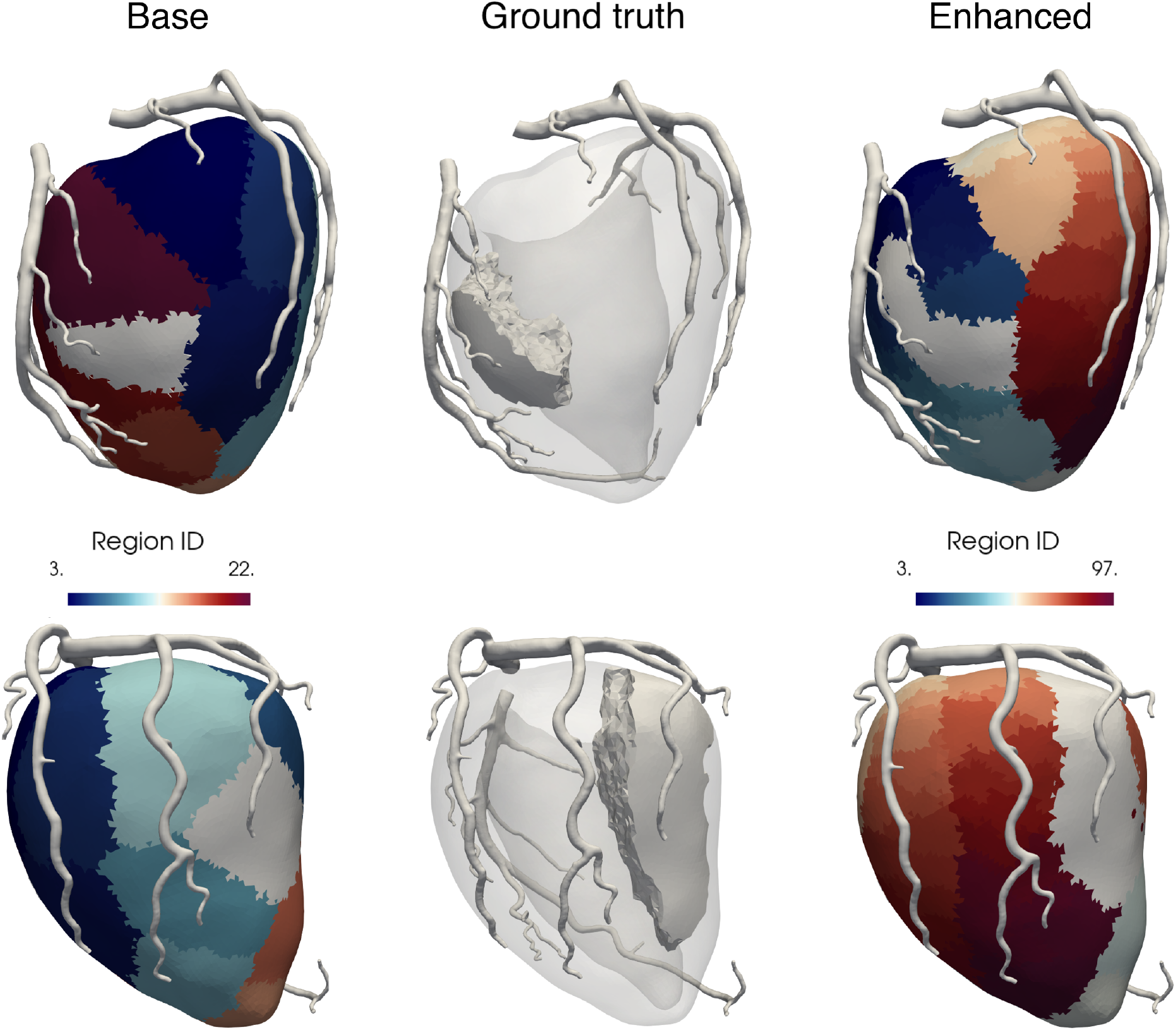
Results of the two subdivision strategies in the anterior and posterior views: each distinct colour represents a single perfusion region associated with a coronary outflow. In grey colour is highlighted the union of the regions associated with branch *d*_2_ (top) and *m*_4_ (bottom), as compared with the *ground truth* regions associated to those branches. *Base* strategy: Voronoi tessellation only from the outflows in the 3D segmentation, projected on the myocardial surface. *Enhanced* strategy: Voronoi tessellation from the projected points of the *base* strategy, plus the additional outflows along the epicardial branches (see Figure 4b).

The presence of transverse outflows along the branches allows to discriminate to a much higher detail whether a portion in the myocardium is perfused by the proximal rather than medial or distal tract of its feeding artery. Figure 6 reports a direct comparison between the myocardial mass attributed to the different tracts of the two largest branches (*m*_2_, *m*_3_) with the two strategies and the *ground truth* subdivision. For both the *base* and the *enhanced* strategies, each single perfusion region is assigned to a specific subset (e.g. in the case of branch *m*_2_: proximal, mid-proximal, mid-distal, distal) if the corresponding coronary outflow lies within the 3.5 cm subdivision used for the *ground truth* subdivision, as discussed in Section 3.1.

**Figure 6:**
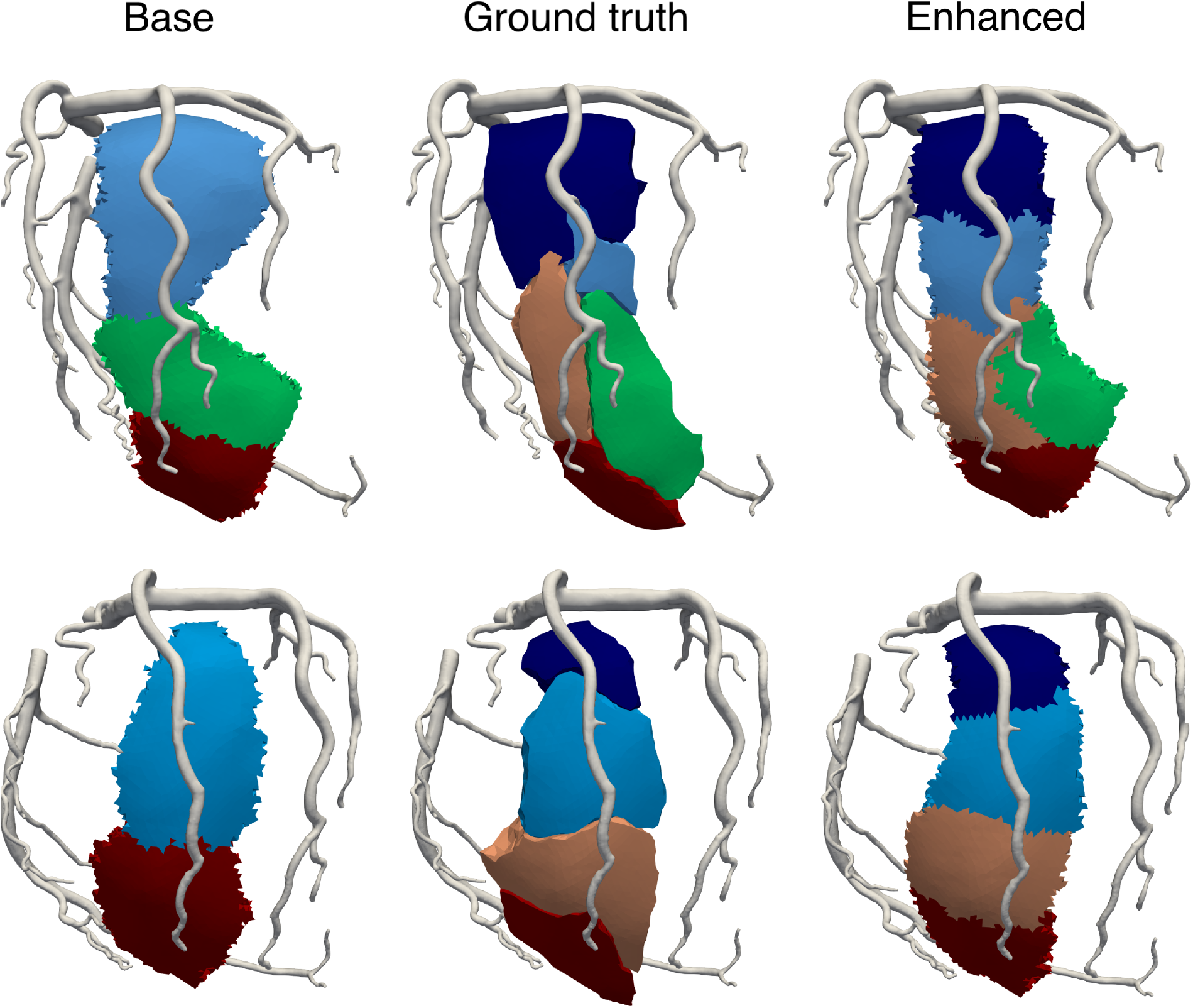
Myocardial mass attribution with the *base* and *enhanced* strategies compared with the *ground truth* perfusion regions for branches *m*_3_ (top) and *m*_2_ (bottom). Each colour represents the myocardial mass attributed to a different subset of the same branch, as explained in Section 3.1.

As we can see, the *enhanced* strategy is in better qualitative agreement with the *ground truth* subdivision and allows to discriminate to a much higher extent between different subsets. In particular, the *base* strategy fails to correctly assign the most proximal myocardial regions, that are erroneously attributed to more distal outflows of the same branch. Also, in the case of bifurcations at the epicardial level the *base* strategy may introduce significant attribution mistakes: this is the case of branch *m*_3_, where most of the mass perfused by one of the sub-branches (distal to the bifurcation) is mistakenly attributed to the other. All these attribution mistakes are significantly reduced using the *enhanced* strategy.

For a quantitative comparison of the two strategies, we compare the *ground truth* perfusion regions with the myocardial subdivision resulting from each strategy and we compute for each branch the amount of misassociated mass following two criteria:

1. Inter-branch misassociation: mass in the *ground truth* perfusion region attributed to the wrong branch;
2. Intra-branch misassociation: mass in the *ground truth* perfusion region attributed to the wrong portion (e.g. proximal, medial, distal) of the correct branch.

We then compute the percentage of misassociated mass dividing it by the total mass of the corresponding *ground truth* perfusion regions. Figure 7 reports the colour-coded representation of the myocardial mass perfused by the LAD alongside the bar plot quantifying the total misassociated mass for the *base* strategy and the *enhanced* strategy using two different values of branching rate: 1.4 and 1 cm^−1^. We report a total misassociated mass (inter-branch and intra-branch) of 24.3% − 48.5% for the *base* strategy, vs 13.1% − 22.8% and 13.7% − 27.5% for the *enhanced* strategy using branching rates of 1.4 and 1 cm^−1^, respectively.

**Figure 7:**
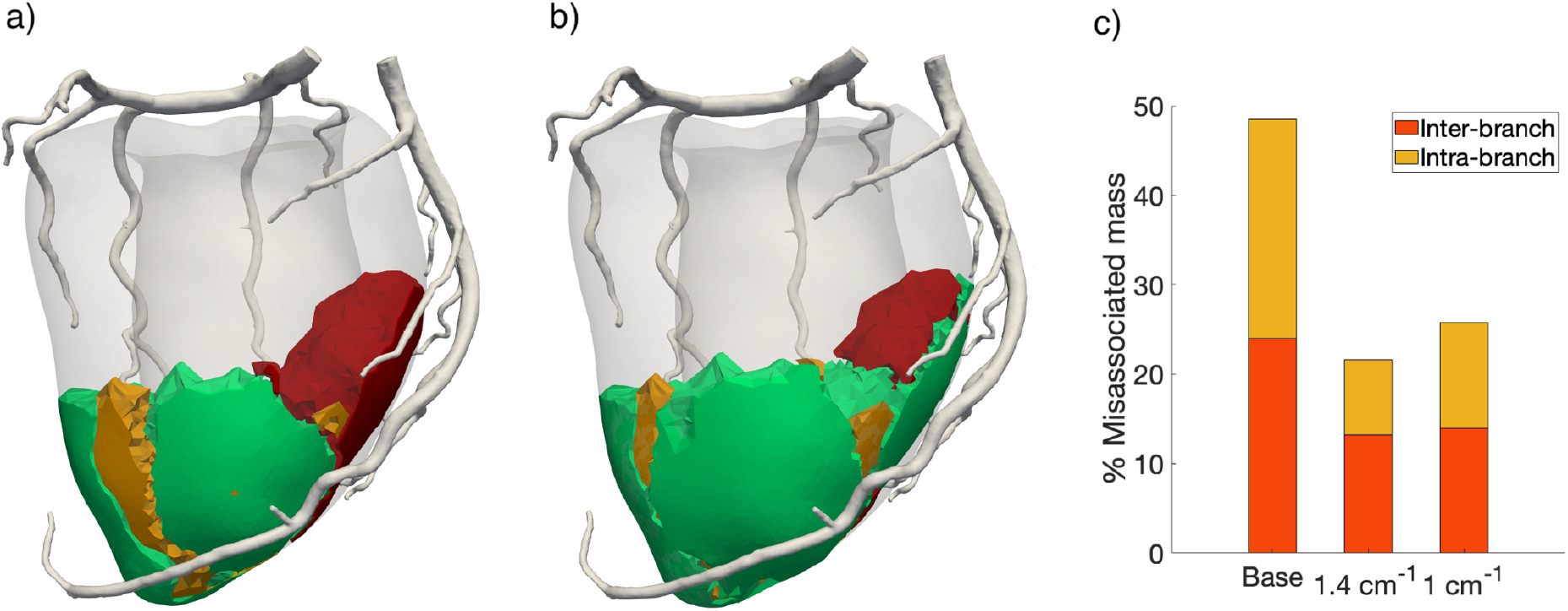
Colour-coded myocardial mass association resulting from the *base* (a) and the *enhanced* (b) subdivision strategies: green represents correctly associated mass, yellow identifies mass with intra-branch misassociation, red identifies mass with inter-branch misassociation. c) Bar plot reporting the total percentage of misassociated mass in the case of the *base* and the *enhanced* strategy using two different branching rates: 1.4 and 1 cm^−1^.

These results indicate that both the inter-branch and, most of all, the intra-branch misassociations decrease significantly when using the *enhanced* strategy with respect to the *base*, showing that the the method we propose leads to a far better accuracy in the coronary-myocardium association. Also, our method allows much higher-precision discrimination between myocardial regions which are perfused by the proximal rather than distal section of each epicardial artery.

This better outcome also shows robustness with respect to the chosen branching rate: we observe no significant increase in inter-branch misassociations (which are the most problematic) and only a slight increase in intra-branch misassociations. This is expected when reducing the number of transverse vessels, since the level of detail that can be achieved when using a coarser branching rate is lower.

## 4. Cardiac perfusion simulations

To explore the impact of the additional transverse outflows, generated by the *enhanced* method, on the haemodynamics in the whole coronary circulation, we run computational simulations of coronary blood flow and cardiac perfusion through a 3D, multiscale finite element model. In Section 4.1 we report the relevant details of the computational model and numerical methods employed, while in Section 4.2 we report the simulations results.

### 4.1. Perfusion simulations: methods

Cardiac perfusion simulations are run using the multiscale computational model we presented in [16]. This model features a 3D fluid dynamics description (Navier-Stokes) for the haemodynamics in the large arteries coupled with a multicompartment Darcy formulation for blood perfusion at the tissue level, featuring a surrogate compliance term which allows for the systolic impediment effect due to cardiac contraction. The coupling between Navier-Stokes and Darcy models is achieved through interface conditions representing mass conservation and balance of interface forces, and it is imposed in such a way that each outflow in the coronary tree is coupled with strictly one perfusion region in the myocardium. All the physics-related parameters of the model (e.g. regarding microvasculature density) as well as the numerical parameters (e.g. space discretization, time step, etc.) are taken from [16]. The inflow boundary condition is a pressure waveform condition (Neumann condition) prescribed at the coronary inlets (representing the aortic pressure curve over time) and is derived following the methods in [13], where we use patient-specific parameters of age and sex that are consistent with the geometry at our disposal (84-year-old healthy female) and we assume normal brachial pressure values of 120-80 mmHg. Since the parameters in [16] were calibrated using hyperaemic flow measures, the scenario simulated in this study represents the haemodynamics in a condition of maximal hyperaemia.

Two simulations are performed to assess haemodynamics in the presence and absence of the additional transverse outflows along the main epicardial branches:

1. *Base* simulation: the coronary tree where the Navier-Stokes problem is solved is the one obtained straight from the 3D segmentation. In this case, the myocardium is subdivided following the *base* strategy discussed in Section 3.1;
2. *Enhanced* simulation: the coronary tree is provided with additional transverse outflows along the main epicardial branches and the myocardium is subdivided following the *enhanced* strategy proposed in Section 3.1.

All the physics-related and numerical parameters are kept the same between the two simulations. For the *enhanced* simulation, the coronary tree used is therefore the same as in the *base* case, with the addition of the transverse outflows placed directly on the coronary wall. Since the mathematical coupling condition imposed at each outflow sets the tangential velocity to zero, this is equivalent to the presence of flow extensions in a direction perpendicular to the wall. This is supported by the vascular data showing that the majority of the transverse vessels branch out in a perpendicular direction (see Figure 2i). In addition, our modelling choices allow for a fully dynamic analysis in a pulsatile regime, which allows us to evaluate the effects of the proposed strategy on local haemodynamics, including blood velocity direction and eventual flow disturbances in the proximity of the additional outflows.

Given that there is no clear pattern for the diameter of the transverse vessels (see Figure 2h), we set all the transverse outflows to have a constant diameter *d* = 0.6 mm for a first analysis. Also, we use a branching rate of 1 cm^−1^; this is done to keep the number of perfusion regions at a manageable level, noticing the robustness of the coronary-myocardium association method we proposed, as discussed in Section 2.2, Figure 7c.

All simulations are run using the software life^x^, a high performance library for Finite Elements simulations of multiphysics, multiscale and multidomain problems developed at MOX - Dipartimento di Matematica, in cooperation with LaBS - Dipartimento di Chimica, Materiali e Ingegneria Chimica, both at Politecnico di Milano [22, 23].

### 4.2. Perfusion simulations: results

Figure 8 reports the numerical results of the *enhanced* simulation, which are consistent with what we obtained in [16] using the *base* strategy on CT-derived geometries. In particular, we can observe the systolic impediment effect, with higher diastolic rather than systolic flow at the capillary level, a reversed pressure gradient in the epicardial arteries at the systolic onset and values of hyperaemic capillary flow between 1 and 6 ml/min/g. These results indicate that the additional transverse outflows do not significantly affect global haemodynamics patterns, which were already well predicted by the model.

**Figure 8:**
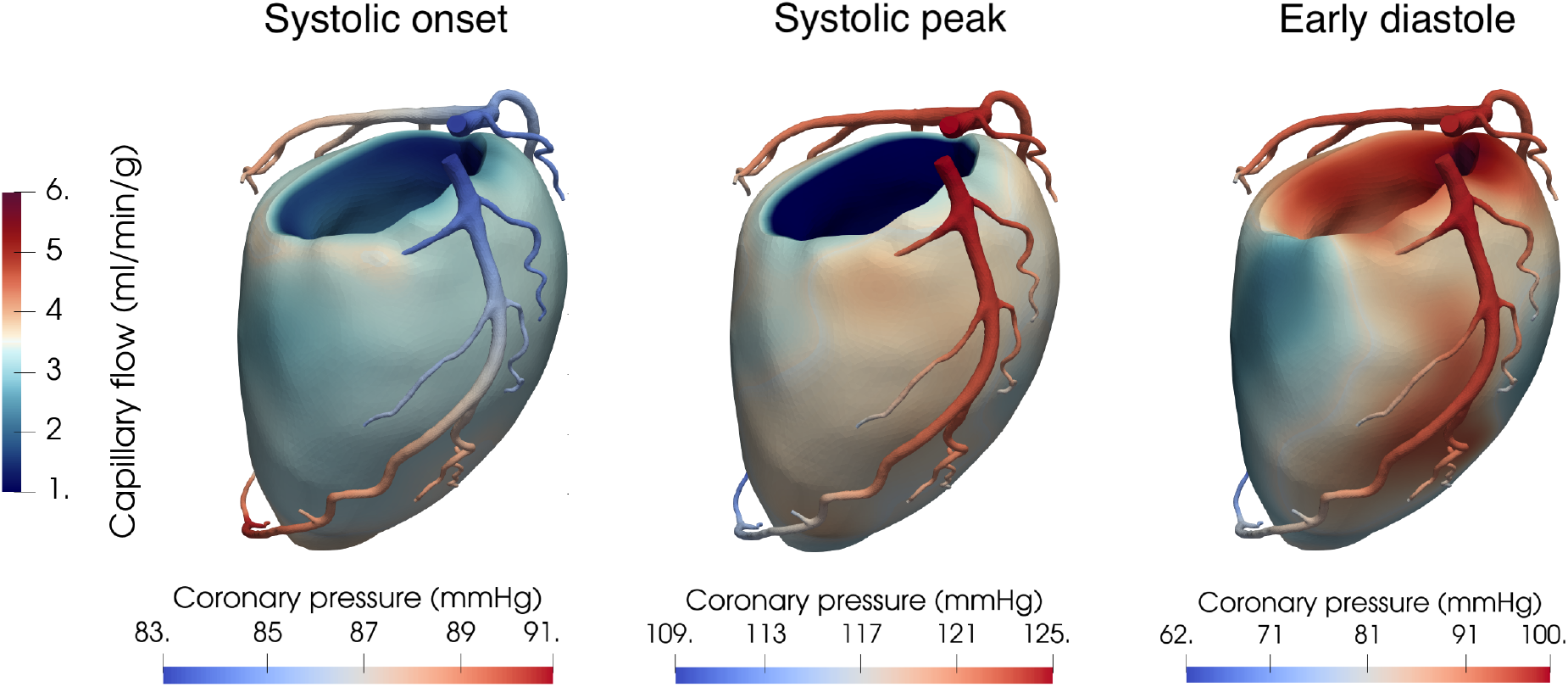
Haemodynamic numerical results for the *enhanced* simulation at three key time instants during the cardiac cycle. The myocardium is colour-coded with the capillary blood flow, whereas in the epicardial coronaries solouring indicates the blood pressure. Notice that the coronary pressure has different scales to better highlight the characteristic features.

Figure 9 reports a detail of the epicardial blood haemodynamics in the proximity of one of the additional transverse outflows. From the super-imposed streamlines depicting the direction of blood velocity we can see that the velocity vectors are perpendicular to the outflow surface during the whole cardiac cycle: this is a consequence of the interface condition imposed at these outflows (see [12] for its definition). We can also see that the presence of transverse outflows does not cause any disturbance in flow at any point of the heartbeat. These results indicate that the addition of outflows directly on the wall surface of the coronaries is effective in modelling transverse vessels even without the presence of a flow extension in the mesh representing the vessel itself.

**Figure 9:**
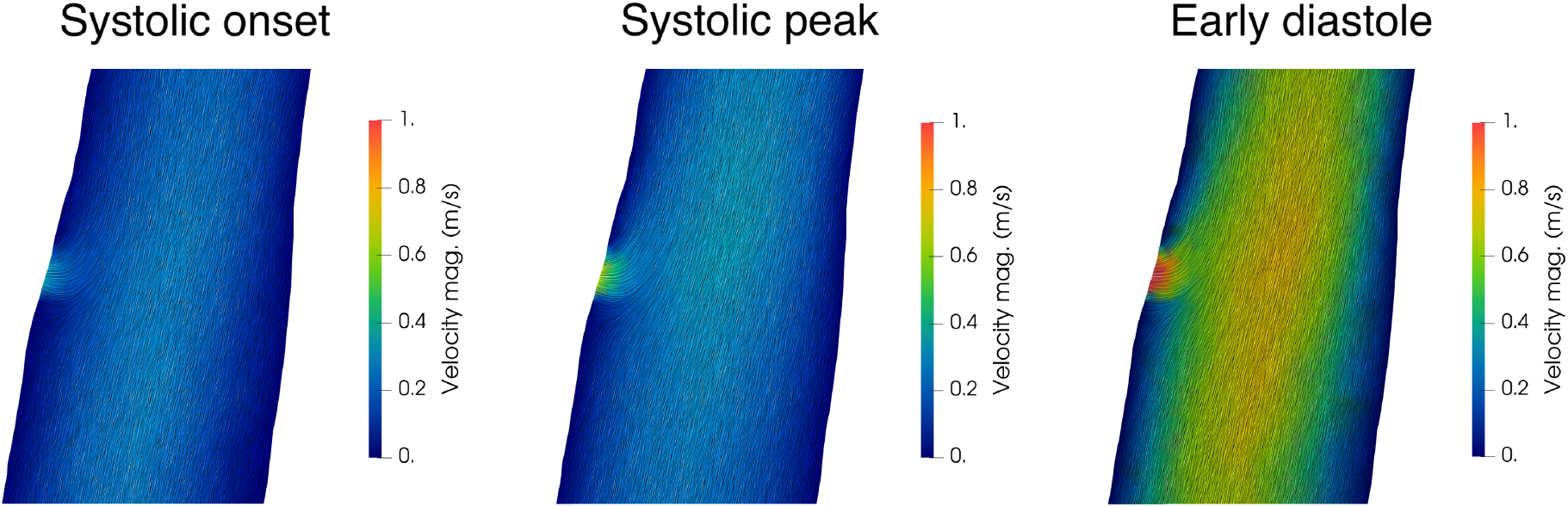
Detail of the coronary blood velocity in the proximity of one of the transverse outflows. On the colour map representing velocity magnitude are super-imposed the streamlines representing velocity direction.

To quantify the effects of the transverse outflows on haemodynamics, we report in Figure 10 a comparison between the predictions of the *base* and *enhanced* simulations regarding clinically useful quantities. We report the profile of the time-averaged coronary pressure along the epicardial arteries, together with the map of Myocardial Blood Flow (MBF) in the myocardium, which is a quantitative measure of the distribution of blood perfusion across the myocardial tissue and is computed as the time-averaged blood flow from the arterioles to the capillaries (see [16] for its mathematical expression).

**Figure 10:**
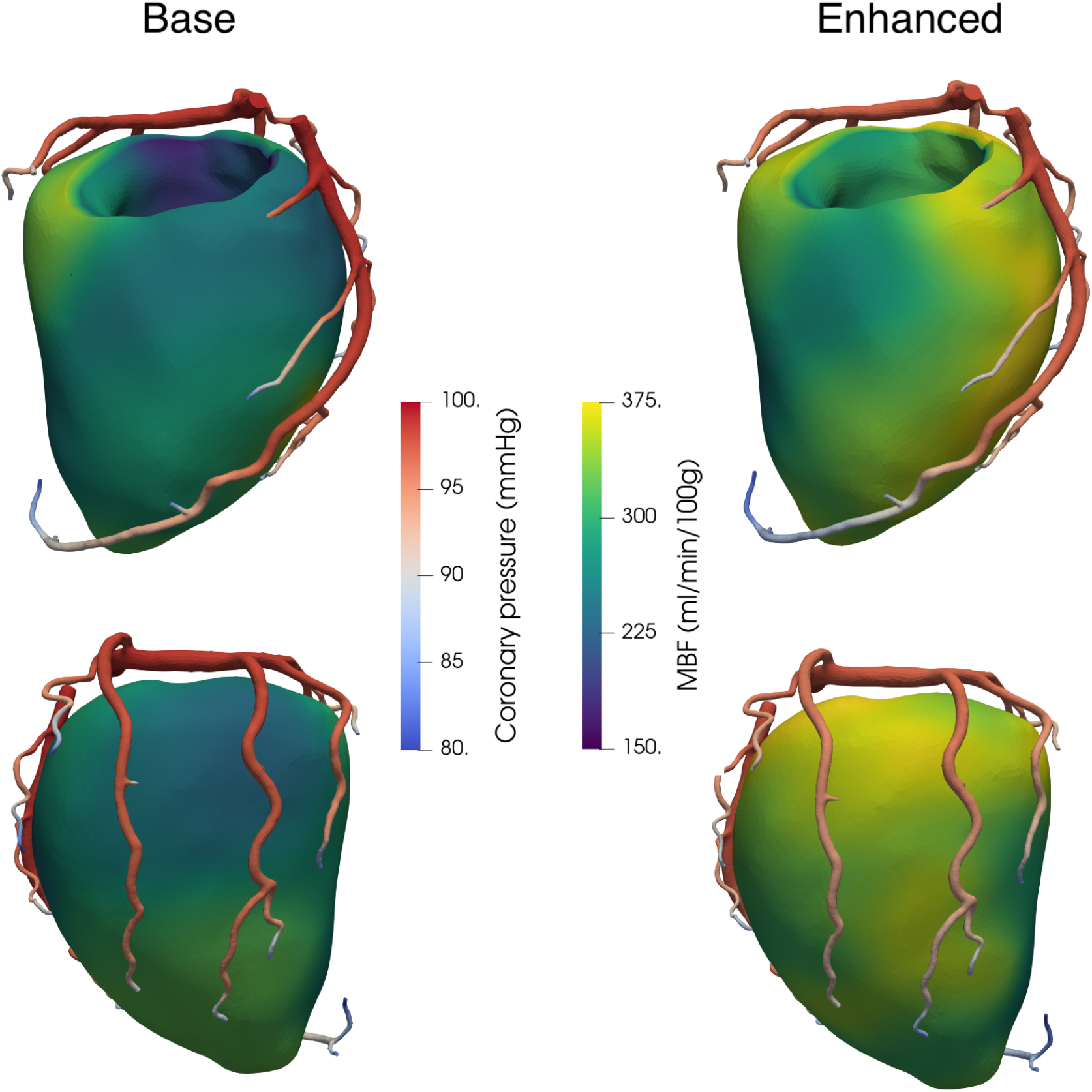
Haemodynamic results for the *base* and *enhanced* simulations. In the epicardial coronaries colouring shows is displayed the profile of the time-averaged coronary pressure, whereas on the myocardium is displayed the Myocardial Blood Flow (MBF) map.

As can be seen in Figure 10, both simulations produce realistic results: pressure gradients along the main coronaries show limited pressure drops (≃ 10 mmHg) with more relevant pressure drops occurring only towards the distal ends of the smaller branches. Regarding MBF, we report an volume average value of 267.5 ml/min/100g and 314.2 ml/min/100g for the *base* and *enhanced* simulations, respectively. Both of these values are consistent with hyperaemic global MBF values for nonpathological individuals [24, 25], although we notice a substantial increase in flow in the *enhanced* simulation: global MBF is 17.4% higher in this case, meaning that a coronary tree which includes additional transverse outflows is less resistive. This is a consequence of the tree structure of the coronaries, where each additional transverse outflow is equivalent to an additional resistance placed in parallel, thus overall reducing the total resistance of the system. This, however, appears to produce negligible effects on the pressure in the epicardial coronaries.

Another major effect is observed in the distribution of MBF throughout the myocardium: in the *base* case, the most perfused regions are typically located at the distal ends of the epicardial branches. This is due to the lack of intermediate outflows, leading to the whole blood flow being carried to the distal ends, the only exceptions being the marginal branches *m*_2_ and *m*_3_. However, also in these cases the presence of a single transverse vessel is not enough to achieve uniform perfusion. In the *enhanced* case, instead, we observe a redistribution of blood flow towards the more proximal regions, with the distal regions being almost unaffected. This more homogeneous perfusion is in much better agreement with previous observations of blood flow across the myocardium in animals and humans [26, 27]. Overall, these results demonstrate that the inclusion of anatomically accurate coupling between coronaries and myocardial tissue has a relevant impact for the distribution of cardiac perfusion and is definitely a factor to take into account in this type of studies.

## 5. Discussion

Impairments in cardiac perfusion represent one of the most relevant clinical challenges of modern times, leading to major adverse events that are responsible for around 17.8 million deaths worldwide each year [28]. This is also due to the extreme complexity in the clinical management of these patients, which includes diagnosis, risk stratification and choice of the best treatment option. In this context, a crucial issue is that the current resolution limit of *in vivo* medical imaging techniques allows the visualization of the major coronary arteries only, introducing uncertainties in the association between the coronary feeding arteries and the perfused myocardial regions. This leads to several clinical issues, for example the correct quantification of the muscle mass perfused by each epicardial branch, the evaluation of which lesions affect a sufficiently large mass to pose a significant threat to the patient, and even the prediction of the outcomes of possible revascularization treatments.

Detailed morphometric data of the entire arterial coronary circulation, available for pigs [1, 2, 3] and recently humans [4], has led to significant insights; however these datasets are highly subject-specific and their direct application to other hearts is limited by the high inter-subject variability in the coronary anatomy, in particular at the epicardial level. Therefore, there is the need for novel strategies to transfer the subject-specific information on coronary morphometry to any given coronary tree, for example reconstructed from minimally invasive medical images such as CT or MRI scans.

The present work aims to fulfill this need, by exploiting quantitative information on coronary branching patterns for the development of a novel tool that can be applied to any epicardial tree to achieve a precise and personalized association between feeding arteries and myocardial mass. Compared to previous work in this direction [7], this study is the first to provide a subdivision strategy with a quantitative validation against detailed morphometric human data. Our results demonstrate that the inclusion of transverse vessels, along the major coronary arteries and their secondary branches, is essential to achieve a correct artery-myocardium association, reducing the amount of misclassified myocardial mass by more than half and showing robustness for different values of branching rate.

The tool we propose can be easily integrated into 3D computational models of coronary and myocardial blood flow, that represent a very promising way to predict the ischaemic burden of coronary artery disease in a noninvasive way. An inherent complexity of these models is their need to provide a spatial link between the major coronaries and the different regions of the myocardium to provide reliable estimates of the distribution of blood flow. This spatial link has been implemented in previous computational studies through Voronoi tessellation of the myocardium starting from the geometry of the coronary arteries. The transverse vessels, however, have been either excluded [16, 14] or generated artificially in a strictly 1D setting [15]. In this work, we provide a way to include transverse vessels in models featuring a 3D description of epicardial haemodynamics, demonstrating that these vessels can be modeled through the simple addition of outflows on the sidewalls of the coronary mesh with a specific choice for the interface condition (constraining to zero the tangential velocity). Our results show that this choice does not impair the local haemodynamics and provide for the first time an effective and efficient way to include transverse vessels in a 3D fluid-dynamics framework.

From a clinical perspective, we report that the inclusion of transverse outflows has a negligible effect on the pressure in the epicardial arteries, thus it is unlikely to influence any related metrics used in the functional evaluation of coronary stenoses, e.g. the Fractional Flow Reserve (FFR) index [29]. On the other hand, there is a relevant effect on the global MBF and its distribution across the myocardium: we observe a redistribution of MBF towards the more proximal regions as well as an overall increase in flow due to the coronary tree becoming less resistive. This indication is particularly relevant given the high variability of the different segmentation methods currently employed for the coronary arteries; the consequence is that the same model can lead to substantially different results depending on the number of secondary branches that are included. A more comprehensive analysis of this aspect is needed to understand whether the presence of fewer visible branches should be taken as an indication of a lower myocardium vascularization or if computational models should adjust their parameters to offset such anatomical differences.

We believe that the methods proposed in this work are a significant step toward a more effective use of medical imaging exams, with some relevant perspective applications discussed in what follows.

A first possible clinical application is the quantification of the Mass at Risk (MaR) index associated with an epicardial coronary lesion. This index quantifies the total muscle mass affected by a specific lesion and is strictly associated with the mass perfused by the portions of the diseased branch located distally to such a lesion. Correct quantification of MaR is clinically relevant, as it has been shown to be strongly correlated with ischaemic burden and to be an independent predictor of major adverse events [5, 30, 9]. However, established methods for MaR quantification have shown sufficient reliability only when based on invasive procedures, such as during forced coronary occlusion in a catheterized setting [31, 32]. Noninvasive MaR quantification based on CT scans has been recently proposed [8], showing good agreement with invasive angiographic estimates at the level of the three major arteries (LAD, LCX, RCA). In the present work, we push forward this idea by associating also secondary branches (e.g. diagonal and marginal branches) to separate perfusion regions, while retaining a hierarchical distinction between proximal, medial and distal portions of each branch. This could allow a more precise MaR quantification in the case of complex lesions affecting the secondary coronary branches rather than the main arteries.

A second application focuses on the use of fully personalized models as digital twins to predict the outcomes of percutaneous coronary intervention (PCI). The presence of complex lesions, featuring diffuse coronary artery disease and multiple stenotic sites, complicates the definition of an optimal revascularization strategy and is associated with a worse outcome [33, 34]. Other works have been successful in predicting post-intervention FFR based on pre-intervention angiographic [35] or CT [36] imaging; however, no study yet has investigated the PCI outcomes in terms of optimal reperfusion of the myocardium. To this aim, a very promising direction is the use of pre-intervention functional imaging (such as dynamic stress CT perfusion or Positron Emission Tomography) to calibrate a digital twin which is later used to make predictions on the best PCI strategy to achieve optimal reperfusion. This strategy can be successful only if the model calibration is performed based on the right association between epicardial branches and myocardial regions, so the tools proposed in this work represent a crucial step to the viability of this application. Also, the inclusion of discrete outflows in a 3D coronary geometry allows interesting investigations on the impact of complex morphologies of the atherosclerotic plaques on how effectively the transverse vessels perfuse the underlying myocardial mass.

This work has some limitations:

1. We validate our method on a dataset of a single heart, so we cannot assess how inter-subject variability in coronary anatomy affects the performances of the proposed strategy. However, since the most relevant anatomical differences regard the course of the major epicardial branches and these are captured also by *in vivo* medical images, we believe that our method retains its accuracy also when applied to other coronary trees. Access to more detailed human datasets would allow a more extensive validation and is, thus, highly desirable;
2. We do not include a validation of the haemodynamic predictions on blood flow distribution across the myocardium against subject-specific measures. However, our results are consistent with literature data for human subjects without obstructive coronary artery disease. Specifically, previous works reported homogeneous blood flow distribution across the myocardium [26, 27] which we report in the case of the coronary arteries that include the additional transverse outflows. We believe this evidence to be sufficient to conclude that our method leads to a better description of coronary haemodynamics at the tissue level.

## 6. Acknowledgements

GMP has been supported by Bracco Imaging S.p.A., by Consiglio Nazionale delle Ricerche (CNR) and by Italian PNRR research funding, Missione 4, DM226/2021.

GMP, CV are members of the INdAM group GNCS “Gruppo Nazionale per il Calcolo Scientifico” (National Group for Scientific Computing).

CV has been partially supported by: i) the European Union-Next Generation EU, Mission 4, Component 1, CUP: D53D23018770001, under the research project MIUR PRIN22-PNRR n.P20223KSS2, “Machine learning for fluid structure interaction in cardiovascular problems: efficient solutions, model reduction, inverse problems”, ii) the Italian Ministry of Health within the PNC PROGETTO HUB LIFE SCIENCE - DIAGNOSTICA AVANZATA (HLS-DA) “INNOVA”, PNCE3-2022-23683266–CUP: D43C22004930001, within the “Piano Nazionale Complementare Ecosistema Innovativo della Salute” - Codice univoco investimento: PNCE3-2022-23683266; iii) the Italian research project MIUR PRIN22 n.2022L3JC5T “Predicting the outcome of endovascular repair for thoracic aortic aneurysms: analysis of fluid dynamic modelling in different anatomical settings and clinical validation”; iv) Italian Ministry of Health within the project “CAL.HUB.RIA” - CALABRIA HUB PER RICERCA INNOVATIVA ED AVANZATA. Code: T4-AN-09, CUP: F63C22000530001.

NAH and JAM gratefully acknowledge the support of the the U.K. EPSRC grant number EP/S030875/1 (SofT-Mech with MIT and POLIMI (SofTMechMP)).

We acknowledge the CINECA award under the ISCRA initiative, for the availability of high performance computing resources and support, in the ISCRA-C project Vr-PCI.

## Footnotes

1 https://www.mathworks.com/help/matlab/ref/alphashape.html

## References

[1] E. VanBavel, J. A. Spaan, Branching patterns in the porcine coronary arterial tree. estimation of flow heterogeneity., Circulation Research 71 (5) (1992) 1200–1212. doi:10.1161/01.RES.71.5.1200. URL https://www.ahajournals.org/doi/abs/10.1161/01.RES.71.5.1200

[2] G. S. Kassab, C. A. Rider, N. J. Tang, Y. C. Fung, Morphometry of pig coronary arterial trees, American Journal of Physiology-Heart and Circulatory Physiology 265 (1) (1993) H350–H365. doi:10.1152/ajpheart.1993.265.1.H350.

[3] B. Kaimovitz, Y. Lanir, G. Kassab, A full 3-d reconstruction of the entire porcine coronary vasculature, American journal of physiology. Heart and circulatory physiology 299 (2010) H1064–76. doi:10.1152/ajpheart.00151.2010.

[4] J. Schwarz, M. van Lier, J. van den Wijngaard, M. Siebes, E. VanBavel, Topologic and hemodynamic characteristics of the human coronary arterial circulation, Frontiers in Physiology 10 (2020) 1611. doi:10.3389/fphys.2019.01611.

[5] M. M. Graham, P. D. Faris, W. A. Ghali, P. Galbraith, C. M. Norris, J. T. Badry, L. Mitchell, M. J. Curtis, M. L. Knudtson, Validation of three myocardial jeopardy scores in a population-based cardiac catheterization cohort, American Heart Journal 142 (2) (2001) 254–262. doi:10.1067/mhj.2001.116481.

[6] M. Termeer, J. Bescós, M. Breeuwer, A. Vilanova, F. Gerritsen, E. Gröller, E. Nagel, Visualization of myocardial perfusion derived from coronary anatomy, IEEE transactions on visualization and computer graphics 14 (2008) 1595–602. doi:10.1109/TVCG.2008.180.

[7] M. Termeer, J. O. Bescós, M. Breeuwer, A. Vilanova, F. Gerritsen, Patient-specific mappings between myocardial and coronary anatomy (2010).

[8] S. Ide, S. Sumitsuji, O. Yamaguchi, Y. Sakata, Cardiac computed tomography-derived myocardial mass at risk using the voronoi-based segmentation algorithm: A histological validation study, Journal of Cardiovascular Computed Tomography 11 (3) (2017) 179–182. doi:10.1016/j.jcct.2017.04.007.

[9] F. van Driest, C. Bijns, R. van der Geest, A. Broersen, J. Dijkstra, J. Jukema, A. Scholte, Correlation between quantification of myocardial area at risk and ischemic burden at cardiac computed tomography, European Journal of Radiology Open 9 (2022) 100417. doi:10.1016/j.ejro.2022.100417.

[10] A. Cookson, J. Lee, C. Michler, R. Chabiniok, E. Hyde, D. Nordsletten, M. Sinclair, M. Siebes, N. Smith, A novel porous mechanical framework for modelling the interaction between coronary perfusion and myocardial mechanics, Journal of biomechanics 45 (5) (2012) 850–855.

[11] J. Lee, A. Cookson, R. Chabiniok, S. Rivolo, E. Hyde, M. Sinclair, C. Michler, T. Sochi, N. Smith, Multiscale modelling of cardiac perfusion, Modeling the heart and the circulatory system (2015) 51–96.

[12] S. Di Gregorio, C. Vergara, G. Montino Pelagi, A. Baggiano, P. Zunino, M. Guglielmo, L. Fusini, G. Muscogiuri, A. Rossi, M. G. Rabbat, A. Quarteroni, G. Pontone, Prediction of myocardial blood flow under stress conditions by means of a computational model, European Journal of Nuclear Medicine and Molecular Imaging 49 (6) (2022) 1894 – 1905. doi:10.1007/s00259-021-05667-8.

[13] G. Montino Pelagi, A. Baggiano, F. Regazzoni, L. Fusini, M. Alì, G. Pontone, G. Valbusa, C. Vergara, Personalized pressure conditions and calibration for a predictive computational model of coronary and myocardial blood flow., Annals of biomedical engineering. URL https://api.semanticscholar.org/CorpusID:267572450

[14] K. Menon, M. Khan, Z. Sexton, J. Richter, P. Nguyen, S. Malik, J. Boyd, K. Nieman, A. Marsden, Personalized coronary and myocardial blood flow models incorporating ct perfusion imaging and synthetic vascular trees, npj Imaging 2. doi:10.1038/s44303-024-00014-6.

[15] L. Papamanolis, H. J. Kim, C. Jaquet, M. Sinclair, M. Schaap, I. Danad, P. van Diemen, P. Knaapen, L. Najman, H. Talbot, C. A. Taylor, Vignon-Clementel, Myocardial Perfusion Simulation for Coronary Artery Disease: A Coupled Patient-Specific Multiscale Model, Annals of Biomedical Engineering 49 (2021) 1432–1447. doi:10.1007/s10439-020-02681-z. URL https://hal.archives-ouvertes.fr/hal-03036457

[16] G. Montino Pelagi, F. Regazzoni, J. M. Huyghe, A. Baggiano, M. Alì, S. Bertoluzza, G. Valbusa, G. Pontone, C. Vergara, Modeling cardiac microcirculation for the simulation of coronary flow and 3d myocardial perfusion, Biomechanics and Modeling in Mechanobiology (2024) 1–26.

[17] J. A. Spaan, R. ter Wee, J. W. van Teeffelen, G. Streekstra, M. Siebes, C. Kolyva, H. Vink, D. Fokkema, E. VanBavel, Visualisation of intramural coronary vasculature by an imaging cryomicrotome suggests compartmentalisation of myocardial perfusion areas, Medical and Biological Engineering and Computing 43 (2005) 431–435.

[18] M. Zubaid, C. Buller, G. Mancini, Normal angiographic tapering of the coronary arteries., The Canadian journal of cardiology 18 (9) (2002) 973–980.

[19] M. A. Bartololo, A. M. Taylor-LaPole, D. Gandhi, A. Johnson, Y. Li, E. Slack, I. Stevens, Z. Turner, J. D. Weigand, C. Puelz, et al., Computational framework for the generation of one-dimensional vascular models accounting for uncertainty in networks extracted from medical images, ArXiv.

[20] J. S. Choy, G. S. Kassab, Scaling of myocardial mass to flow and morphometry of coronary arteries, Journal of Applied Physiology 104 (5) (2008) 1281–1286. doi:10.1152/japplphysiol.01261.2007.

[21] D. C. J. Keulards, S. Fournier, M. van ‘t Veer, I. Colaiori, J. M. Zelis, M. El Farissi, F. M. Zimmermann, C. Collet, B. De Bruyne, N. H. J. Pijls, Computed tomographic myocardial mass compared with invasive myocardial perfusion measurement, Heart 106 (19) (2020) 1489– 1494. doi:10.1136/heartjnl-2020-316689.

[22] P. C. Africa, lifex: A flexible, high performance library for the numerical solution of complex finite element problems, SoftwareX 20 (2022) 101252.

[23] P. C. Africa, I. Fumagalli, M. Bucelli, A. Zingaro, M. Fedele, L. Dede’, A. Quarteroni, lifex-cfd: An open-source computational fluid dynamics solver for cardiovascular applications, Computer Physics Communications 296 (2024) 109039.

[24] K.-T. Ho, H.-Y. Ong, G. Tan, Q.-W. Yong, Dynamic ct myocardial perfusion measurements of resting and hyperaemic blood flow in low-risk subjects with 128-slice dual-source ct, European Heart Journal-Cardiovascular Imaging 16 (3) (2015) 300–306.

[25] L. Lyu, J. Pan, D. Li, X. Li, W. Yang, M. Dong, C. Guo, P. Lin, Y. Han, Y. Liang, et al., Knowledge of hyperemic myocardial blood flow in healthy subjects helps identify myocardial ischemia in patients with coronary artery disease, Frontiers in Cardiovascular Medicine 9 (2022) 817911.

[26] P. Chareonthaitawee, P. A. Kaufmann, O. Rimoldi, P. G. Camici, Heterogeneity of resting and hyperemic myocardial blood flow in healthy humans, Cardiovascular research 50 (1) (2001) 151–161.

[27] E. Y. Kim, W.-J. Chung, Y. M. Sung, S. S. Byun, J. H. Park, J. H. Kim, J. Moon, Normal range and regional heterogeneity of myocardial perfusion in healthy human myocardium: assessment on dynamic perfusion ct using 128-slice dual-source ct, The International Journal of Cardiovascular Imaging 30 (2014) 33–40.

[28] J. C. Brown, T. E. Gerhardt, E. Kwon, Risk factors for coronary artery disease.

[29] P. A. Tonino, B. De Bruyne, N. H. Pijls, U. Siebert, F. Ikeno, M. vant Veer, V. Klauss, G. Manoharan, T. Engstrøm, K. G. Oldroyd, et al., Fractional flow reserve versus angiography for guiding percutaneous coronary intervention, New England Journal of Medicine 360 (3) (2009) 213–224.

[30] K. Sadamatsu, K. Nagaoka, Y. Koga, K. Kagiyama, K. Muramatsu, K. Hironaga, H. Tashiro, T. Ueno, Y. Fukumoto, The functional severity assessment of coronary stenosis using coronary computed tomography angiography-based myocardial mass at risk and minimal lumen diameter, Cardiovascular Therapeutics 2020 (1) (2020) 6716130.

[31] L. Ceriani, E. Verna, L. Giovanella, L. Bianchi, G. Roncari, G. L. Tarolo, Assessment of myocardial area at risk by technetium-99m sestamibi during coronary artery occlusion: comparison between three tomographic methods of quantification, European journal of nuclear medicine 23 (1996) 31–39.

[32] O. Pereztol-Valdés, J. Candell-Riera, G. Oller-Martínez, S. Aguadé-Bruix, J. Castell-Conesa, J. Ángel, J. Soler-Soler, Localization and quantification of myocardium at risk by myocardial perfusion spect during coronary artery occlusion, Revista Española de Cardiología (English Edition) 57 (7) (2004) 635–643.

[33] M. O. Mohamed, J. Polad, D. Hildick-Smith, O. Bizeau, R. K. Baisebenov, M. Roffi, A. Iniguez-Romo, B. Chevalier, C. von Birgelen, A. Roguin, et al., Impact of coronary lesion complexity in percutaneous coronary intervention: one-year outcomes from the large, multicentre e-ultimaster registry, EuroIntervention 16 (7) (2020) 603–612.

[34] J. M. Hanna, S. Y. Wang, A. Kochar, D. Y. Park, A. A. Damluji, G. A. Henry, Y. Ahmad, J. P. Curtis, M. G. Nanna, Complex percutaneous coronary intervention outcomes in older adults, Journal of the American Heart Association 12 (19) (2023) e029057.

[35] R. C. Gosling, P. D. Morris, D. A. Silva Soto, P. V. Lawford, D. R. Hose, J. P. Gunn, Virtual coronary intervention: a treatment planning tool based upon the angiogram, JACC: Cardiovascular Imaging 12 (5) (2019) 865–872.

[36] M. Belmonte, M. Maeng, C. Collet, B. L. Norgaard, H. Otake, B. Ko, B.-K. Koo, T. Mizukami, A. Updegrove, E. Barbato, et al., Accuracy of a virtual pci planner based on coronary ct angiography in calcific lesions, Journal of Cardiovascular Computed Tomography 17 (5) (2023) 367–369.

